# Replicating and cycling stores of information perpetuate life

**DOI:** 10.1101/149294

**Authors:** Antony M. Jose

## Abstract

Life is perpetuated through a single-cell bottleneck between generations in many organisms. Here, I highlight that this cell holds information in two distinct forms: in the linear DNA sequence that is replicated during cell divisions, and in the three-dimensional arrangement of molecules that can change during development but that is recreated at the start of each generation. These two interdependent stores of information – one replicating with each cell division and the other cycling with a period of one generation – coevolve while perpetuating an organism. Unlike the genome, the cycling arrangement of molecules, which could include RNAs, proteins, sugars, lipids, etc., is not well understood. Because this arrangement and the genome are together transmitted from one generation to the next, analysis of both is necessary to understand evolution, origins of inherited diseases, and consequences of genome engineering. Recent developments suggest that tools are in place to examine how all the information to build an organism is encoded within a single cell, and how this cell code is reproduced in every generation.

## Introduction

One of the amazing aspects of living things is that they transmit the information for building themselves from one generation to the next. While much of what an organism is made of is specified by the linear sequence information in its DNA genome, the genome is not the only store of information that is transmitted across generations. We can see evidence for the transmission of extra-genomic information when changes that do not alter DNA sequence, nevertheless persist for many generations. For example, in the ciliate *Paramecium aurelia*, changes in the cortex created using explants could be transmitted for many generations (Beisson and Sonneborn, 1965). Some changes reverted after 30-40 generations, and other changes could apparently be maintained indefinitely. Such inheritance of extra-genomic changes invites a consideration of the scope and formulation of all the inherited information that specifies the developing organism’s ensemble of traits – not only the information encoded in the sequence of bases in the DNA, but also the non-genetic information encoded in molecular assemblies independent of DNA sequence.

The information needed to perpetuate an organism must be present even in the life stage that has the least number of cells. This stage serves as a bottleneck for the transmission of information from one generation to the next and is minimally a single cell. Therefore, all the molecules and their arrangement that is reproduced in every such bottleneck stage of an organism is the minimal information that specifies the perpetuation of that organism. This minimal information includes the genome sequence and a spatial arrangement of the genome along with other pre-existing molecules, including RNAs, proteins, sugars, lipids, etc. Organisms transmit these two stores of information from one generation to the next using different strategies. The linear DNA sequence is transmitted by replicating it at each cell division. This strategy allows for retention throughout development of largely the same sequence (barring random mutations) in every cell that forms a continuous lineage from one generation to the next. The three-dimensional arrangement, on the other hand, is transmitted by cycling it with a period of one generation. This strategy allows for potentially extensive transformation of the arrangement throughout development before returning to a similar configuration in the next generation. In addition to changes in the three-dimensional arrangement of molecules, components could be replenished or added to using the DNA sequence and raw materials from the environment over the course of a life cycle. Thus, life is perpetuated by two stores of information that are interdependent yet distinct.

The need for analyzing both forms of information together is evident if we consider a simple regulatory loop where a transcription factor binds its own promoter and sustains its production in every cell. At the start of each generation, in addition to general cellular machinery, this ‘developmental program’ needs the DNA sequence encoding for the transcription factor, the promoter sequence that the transcription factor binds to, *and* the transcription factor arranged together such that the transcription factor can bind to its binding site in the promoter. In other words, developmental programs are specified using DNA sequence and an arrangement of additional pre-existing molecules (see Dyson 1982, Maynard Smith, 1990, Ganti, 2003, and Ptashne 2013 for related views).

Here, I develop this perspective on inherited information further, discuss its implications, and suggest approaches for the discovery of the non-genetic store of information that is transmitted along with the genome across generations to drive development and evolution. While this article has an emphasis on animal biology to give it focus, the concepts presented here are applicable to all cellular life: bacteria, archaea, and eukarya.

### Organisms cycle minimally through a single cell

The continuity of life relies on cycles. Single-celled organisms go through cell division cycles (Murray and Hunt, 1993). Parasites go through elaborate cycles within different cells and even different host organisms (Olsen, 1986). Multicellular organisms cycle through a single-cell bottleneck – the fertilized zygote in sexually reproducing organisms – that carries all the information necessary to make the next generation (Wilson, 1896). In animals, the zygote can either be present within an egg that is laid or be implanted within the uterus of the mother and grow as a fetus. In animals that lay eggs, all the information necessary for development is contained within the fertilized egg. Under permissive environmental conditions, a hatchling is inevitable. In animals that give birth to live young, the growing fetus shares circulation with the mother, which can be thought of as providing the permissive environment and more. Organisms that undergo parthenogenesis (Phillips, 1903) also cycle through a single-cell stage, an egg that does not need fertilization by sperm. Finally, regenerating organisms (Birnbaum and Sanchez-Alvarado, 2008) can begin each generation from many different collections of cells. Nevertheless, in every scenario, the simplest life stage of an organism minimally consists of a single cell.

### A single-cell bottleneck constrains multicellularity

The ‘beginning’ of an organism’s life is an arbitrary time-point in its life cycle (McLaren, 1980). For multicellular organisms that reproduce sexually, if we consider the zygote generated by the union of gametes to be the beginning, what organisms can this cell develop into while ensuring the continuity of life? An initial answer could be any collection of cells and material that is compatible with the production of the zygote for the next generation. Indeed, multicellular organisms have evolved many different types of cells with very different properties (Arendt et al., 2016) – consider the sciatic nerve that can grow to be a meter long versus a tiny lymphocyte in humans. In most cases the DNA within every cell is essentially the same. The drastic differences between cells are achieved by changing how the same DNA is used to make RNA, protein, and other components within a cell. As a powerful demonstration of this principle, the nucleus of any cell can be combined with the cytoplasm of an oocyte to recreate an entire animal (Briggs and King, 1952; Gurdon et al., 1958) and the addition of a few factors can reprogram one cell type into another (Lassar et al., 1986; Takahashi and Yamanaka, 2006). Notable exceptions to this general rule include human red blood cells, which jettison their nuclei as they mature (Thompson, 1951). Nevertheless, the collection of remarkably different cells that can make up an organism poses no problems for the continuity of life as long as it is compatible with the creation of a zygote to start the next generation, hence the famous saying “a hen is only an egg’s way of making another egg” (Butler, 1910). Expanding these considerations to populations, however, reveals that individual members of a population can be exempt from this constraint as long as they support one individual that is bound by the constraint to produce a zygote. For example, in colonies of bees or ants, drones or workers can be sterile and yet support the queen, which perpetuates the colony (Heinze and Schrempf, 2008). These constraints focus our attention on the cell – the zygote in many cases – as an important unit of organization that needs to be understood to explain life.

### Much of the information needed to specify the zygote of any organism is unknown

Factors that identify a specific zygote include both the contents of the cell and the spatial arrangement of the contents, which can be crucial for function. Imagine a “typical” eukaryotic cell with many typical components like mitochondria, lysosomes, endoplasmic reticulum, and other organelles; ribosomes, polymerases, and other molecular machines; ATP, Ca^2+^, and other small molecules; and so on. Everything in this cell needs to be organized in three dimensions for appropriate function. *What would we need to add to our description of this typical cell to describe the zygote that can develop into a particular organism?* The answer to this question is likely to include many different molecules. For example, a genome with appropriate chromatin to allow access to early transcription factors, mRNAs that are poised to be translated, and sufficient stores of small molecules and enzymes that use the molecules to initiate development in permissive environments. Each of these molecules likely needs to be present at precise amounts as illustrated by the control of mRNA levels in the zygote, which is tuned to allow development (Gerson-Gurwitz et al., 2016). Attempts have been made to comprehensively catalogue different components of cells: the genome, the epigenome, the transcriptome, the proteome, the metabolome, etc. These techniques can be used to discover the contents of a particular zygote. Discovering the spatial arrangement of components within a cell, however, has been more difficult but attempts to generate a comprehensive map of at least the proteins within a few cell types are underway (Thul et al., 2017; Allen Institute for Cell Science). Yet, spatial arrangement and sub-cellular localization are crucial for function. Furthermore, we now know that the genome is organized into spatial domains that impact gene expression in eukaryotes (reviewed in Bonev and Cavalli, 2016). The functions of key molecules within cells are regulated through chemical modifications that continue to be discovered (e.g. RNA modifications (Helm and Motorin, 2017), DNA modifications (Sood et al., 2016), histone modifications (Andrews et al., 2016), etc.). Finally, we currently do not understand how the components within a cell determine many aspects of cellular behavior (see Marshall, 2015 for an interesting list). Thus, there is much to learn before we can specify the molecules and their arrangement at a given time that describes the zygote of any organism.

### DNA proposes, cell disposes

While the genome is a repository of all sequences that can be transcribed and used for making other components, at any given moment, which RNA is transcribed from DNA depends on what else is within the cell and on the cell’s interactions with the external environment. The information contained in the genome of an organism is thus not sufficient to make that organism. To appreciate this insufficiency, consider a single cell in an organism: (a) the DNA within this cell does not encode all aspects of all molecules in the cell; and (b) whether a molecule is made using the DNA depends on other contents in the cell.

*The linear DNA sequence is not sufficient to build the three dimensional cell*: The RNA and protein components of a cell ultimately rely on DNA for production through the process of transcription and translation using smaller molecules, nucleotides and amino acids, as raw materials. These proteins and RNA can go on to catalyze the formation of additional components such as lipids and sugars from other raw materials. However, the three-dimensional structure and spatial arrangement of all the molecules within a cell is not encoded in the linear DNA. For example, where a protein is localized in the cell can change over time based on interactions with other components within a cell (Thanbichler and Shapiro, 2008). Components in organelles such as mitochondria are not entirely encoded by the nuclear genome and can be considered as endosymbionts with partial autonomy within a eukaryotic cell (Mereschkowsky, 1905; Sagan, 1967). While the DNA encodes the sequence of a protein, the structure it takes depends on its immediate environment (Englander and Mayne, 2014). Environmental changes can even induce cells to over-ride defects caused by mutations in their DNA. For example, mutation of the small GTPase Rac1 can invert the polarity of epithelia and yet normal polarity can be restored by the addition of exogenous laminin protein (O’Brien et al., 2001). Finally, the sizes, numbers, and shapes of organelles, and indeed the collective three-dimensional architecture of a cell all depend on dynamic interactions between components within the cell and with its external environment potentially independent of the DNA (Rafelski and Marshall, 2008).

*Many cell types use the same genome*: The presence of different cell types that retain their identity over time and across cell divisions within an organism reveals that the contents of a cell can exert a controlling effect on the DNA. The mechanisms that make and maintain different cell types were initially referred to as epigenetic control systems (Nanney, 1958) to signify that they are above (‘epi’) genetic control. Pioneers studying such control mechanisms in bacteria imagined multiple modes of regulation for components within a cell in the context of biochemical reactions (Monod and Jacob, 1961), many examples of which have been characterized in the past half-century. These regulatory modes provide teleonomic (i.e programmed) constraints that explain “how come?” rather than “what for?” when events happen within a cell (Mayr, 1961). Furthermore, computational exploration suggests that even random networks of elements with high molecular specificity can result in the emergence of different cell states that remain stable over time (Kauffman, 1969). Extensions of these ideas have led to the identification of gene regulatory networks in model organisms (Levine and Davidson, 2005), where collections of active genes and gene products maintain different cell types while being able to respond to signals during development.

### Replicating and cycling information together drive development

Since its beginnings as *Entwickelungsmechanik* more than a century ago (Sander, 1991), the rich field of developmental biology has been addressing various aspects of how an organism is made. But, the minimal information that is necessary to specify the development of a particular organism in a given external environment is still unknown, and is in fact, relatively unexplored.

Consider an organism that progresses from one generation to the next through a single-cell stage, as is the case with humans and most known multicellular organisms. To discover the minimal information that is necessary for the development of this organism, the entire life cycle of the organism needs to be examined (Figure 1A). For chicken, this would mean examining from egg to egg and not from egg to hen (developmental biology) or from hen to egg (reproductive biology). Let us define everything that is produced using the DNA within the unicellular stage – macromolecules plus organelles plus spatial arrangement – as “C” and the interacting aspects of the environment as “E” (Figure 1B). These external inputs include factors that influence (e.g. temperature), interact with (e.g. extracellular signal), or are imported into (e.g. sugars) the cell.

**Figure 1.**
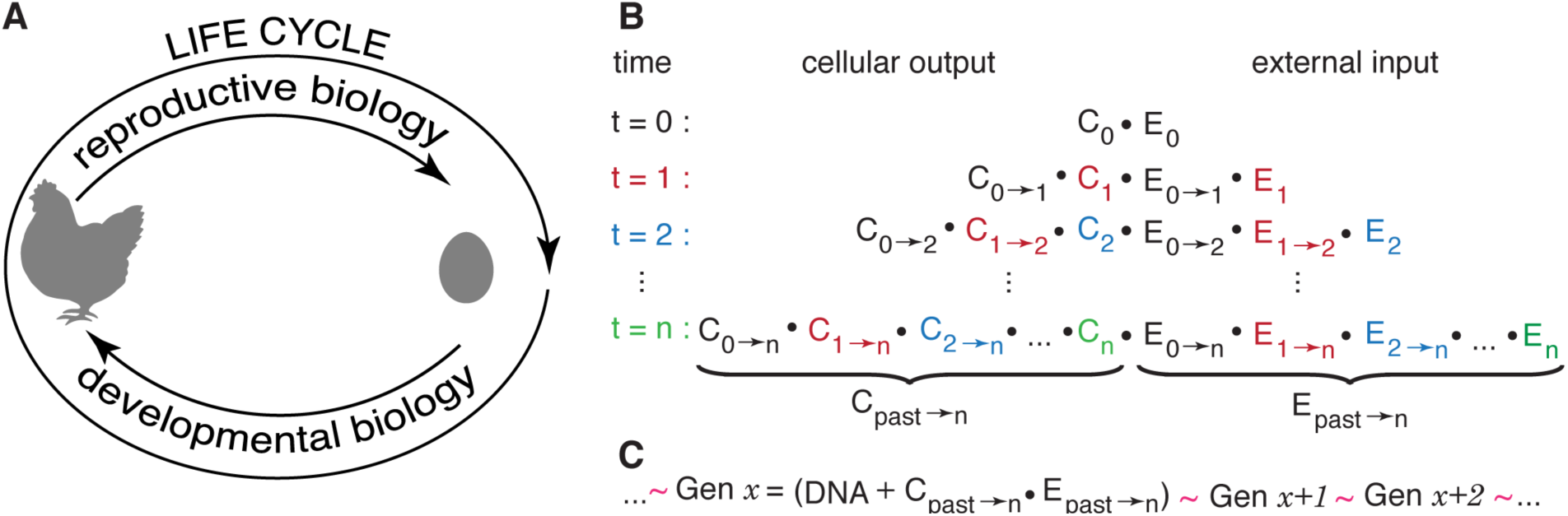
Perpetuation of life. **(A)** Understanding what perpetuates life requires examination of entire life cycles in addition to the study of how organisms are made (developmental biology) and how they reproduce (reproductive biology). (**B**) Every cell accrues components guided by the action of preexisting components on DNA (C_0_, C, C_2_,…). The new components made at each moment (t) can interact (•) with preexisting components, which could have changed since they were made (e.g. C_0>2_ and C_1>2_ at t = 2). This process extends to external components the cell interacts with or acquires independent of DNA within the cell (E_0_, E, E_2_,…). At a given time (t = n), the accrued cellular output (C_pasl>n_) interacts with the accrued external input (E_past>n_). (**C**) DNA and a specific arrangement of components accrued over time that are together required to perpetuate an organism and contained within the single cell that begins each generation (Gen x) defines the cell code of that organism. Recreation of a similar (∼) code in successive generations perpetuates life and is a hallmark of organisms that constrains their evolution.

At an arbitrary initial time, t = 0, C_0_ includes RNA and proteins made using the DNA and other components present in the cell at that time and E_0_ includes molecules in the external environment (including other cells) that is independent of the DNA within the cell. At this arbitrary initial time these components can interact and/or modify each other (depicted as •). As time progresses, everything in the cell is free to change while the DNA is being used to make new RNA and proteins, and more external material can interact with the cell or be imported into the cell. That is, at time t = 1, C_0_ made at t = 0 has changed (C_0_→1) and can now interact with and/or modify (•) C_1_ made at t = 1. Similar changes can also occur in the imported or interacting external material (E_0_→1• E_1_). Furthermore, both the cellular output and the external input can interact and/or modify each other. These changes over time include production of new material (e.g. protein synthesis, lipid synthesis, etc.) and destruction or modification of old material (e.g. autophagy, phosphorylation of lipids, etc.). This process of iterative accumulation of material and progressive change in the state of a cell implies that the DNA *and* everything else accrued in the past that is now within the cell together predicts the next state of a cell in a given environment.

Early attempts at unifying biology focused on understanding the cell in development and inheritance (Wilson, 1896). More than a century ago, some experimental embryologists clearly appreciated the iterative progression (e.g. Boveri, 1902) and influence of history (e.g. Boveri, 1906) when describing the nature of organisms. The interdependence of genes and non-genetic factors within a cell was also well appreciated (e.g., Conklin, 1908; Waddington, 1962). In fact, the term ‘epigenetics’ was initially defined as “…causal mechanisms by which the genes of the genotype bring about phenotypic effects” (Waddington, 1942), but later stated as “the causal interactions between genes and their products which bring the phenotype into being” (Waddington, 1968), making it clear that the genotype (DNA) alone does not bring the phenotype into being. These insights from embryology were also recognized by early proponents of the Modern Synthesis that combined Mendelian genetics with Darwinian evolution (Huxley, 2010), and when ignored, can lead to the mistaken popular view that DNA is the blueprint of life.

In the context of an organism, the contents of a cell can change dramatically over time independent of changes driven by the DNA within the cell. For example, consider progression from one generation to the next in the worm *C. elegans* (Hubbard and Greenstein, 2005) – the best characterized animal we know. From the single-celled zygote, two primordial germ cells are established five cell divisions later. These two cells then go through an extended period of quiescence while the rest of the organism develops. Towards the end of this period intestinal cells cannibalize a large portion of the cytoplasmic contents of both cells (Abdu et al., 2016). Then, the cells proliferate to generate germ cells that subsequently differentiate, first producing sperm, and then oocytes. Maturing oocytes acquire cytoplasmic contents from the rest of the germline (Wolke et al., 2007) and yolk from the intestine (Grant and Hirsh, 1999) potentially along with extracellular RNAs (Marré et al., 2016; Marré and Jose, 2017; Wang and Hunter, 2017). These acquisitions make the oocyte larger than all the 558 cells of a hatching larva combined. Fertilization of this enlarged oocyte creates the zygote for the next generation. Both dramatic change in this cycle – loss of cytoplasmic material from primordial germ cells and gain of cytoplasmic material by oocytes – occur during periods of relative transcriptional quiescence.

Explicitly considering the influence of time and loss/acquisition of non-genetic material, we can depict the single cell that begins each generation at an arbitrary time t = n as DNA plus components accrued until that time and their three-dimensional arrangement i.e. DNA plus C_past_→n•E_past_→n. Everything in this cell that is nearly reproduced in the zygote of successive generations could be necessary to specify the making of the organism. For convenience, hereafter I shall refer to this minimal information that encodes the making of an organism and is contained within a single cell as the cell code for the organism. The upper limit for the cell code is the entire contents and their arrangement in the zygote that begins each generation, i.e., maximally the cell code of the organism = [DNA + C_past_→n•E_past_→n] (Figure 1C). However, the cell code need not equal everything in a zygote. If only a subset of the components and arrangement within a zygote are reproduced in every generation (Figure 2), then the cell code can be less than the contents of a zygote. Furthermore, because many molecular assemblies in life are capable of self-organization (e.g. mitotic spindle (Salmon, 1975)) and templated processes (e.g. transcription (Weiss and Gladstone, 1959)), it is possible that essentially the same cell code could be specified using a subset of the molecules and arrangement that are recreated in every zygote. This possibility is supported by experiments on single-cell regeneration in the ciliate *Stentor polymorphus* - 1/27^th^ of its initial volume can regenerate all structural features (Lillie et al., 1896).

**Figure 2.**
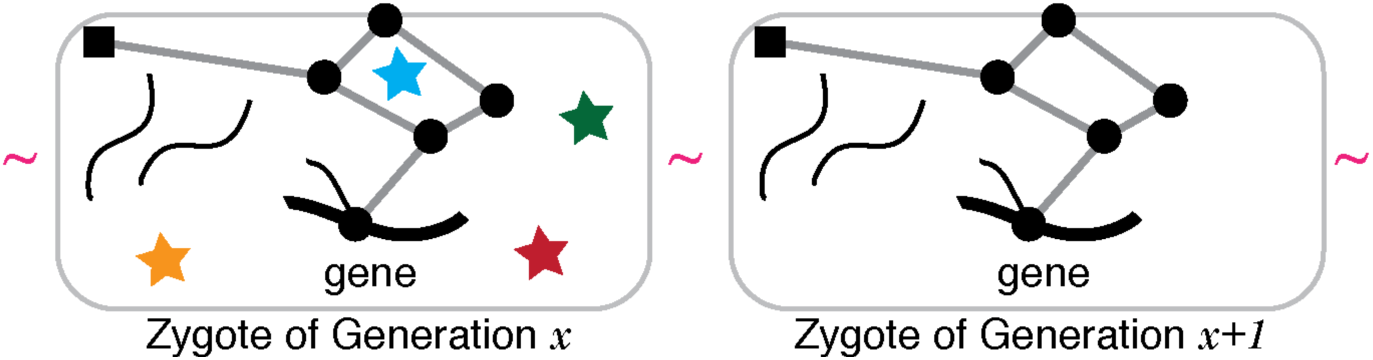
Nature of the cell code. Part of the cell code from the perspective of a single gene (black) and its RNA (curved lines) and protein (square) is depicted. Complexes of other proteins/RNAs/lipids etc. (circles) potentially localized in specific places within the cell and interactions between them (grey lines) that can impact the gene and that are reproduced in every generation are part of the cell code. Some components in the zygote may not be made in successive generations (stars) and compensatory changes in the rest of the zygote may permit different configurations for this gene while preserving the overall cell code of the organism.

Determining the minimal representation of the cell code of an organism requires first determining the reproducible characteristics of zygotes and then answering a set of questions at the molecular level that have rarely, if ever, been posed in experimental systems. For example, imagine a set of characteristics that we have determined about a gene and its products in an organism that are all reproduced in the zygote of every generation, i.e., that are all part of the cell code. Let’s say this gene has chromatin modifications, 2 copies of partially transcribed RNA, 32 copies of the spliced RNA, 17 copies of translated protein, 12 copies of the protein with correct subcellular localization etc. Which of these details are necessary for the perpetuation of a similar arrangement? What changes to the rest of the genome can impact this gene and the arrangement of its products while perpetuating life? What other arrangements of these components are essentially equivalent?

A problem that is separate from determining the cell code of an organism is determining how it is reproduced in every generation. Reproduction of the genetic component of the cell code, i.e. DNA, is relatively trivial and is simply through the process of DNA replication at each cell division. Reproduction of the non-genetic components of the cell code, i.e. the spatial arrangement of molecules including DNA, is likely to be more complex. One possibility is that these arrangements are preserved through cell divisions despite differentiation and development of the organism such that a continuous line of cells from the zygote of one generation to the next carries the cell code for the organism. Alternatively, and more likely, these arrangements vary during differentiation and development but go through an elaborate cycle such that they return to a similar configuration in the zygote of each generation. In the case of unicellular organisms, the cell code needs to be reproduced at a certain time in each cell division cycle. In organisms such as plants where many cells are capable of generating the entire organism, every such cell needs to be able to reproduce the cell code when exposed to conditions that stimulate differentiation and development. Currently, we have a fragmentary understanding of changes in a few potential components of the cell code during development and are far from understanding the mechanism(s) by which the cell code is reproduced in any organism. Nevertheless, a range of components and arrangements are likely to be nearly equivalent cell codes for an organism. We already know that the genomes of individuals within a species can vary a little and yet result in the generation of similar organisms. Additionally discovering alterations to non-genetic aspects of the cell code that are compatible with the perpetuation of life will reveal the full extent of novelty that can arise from one generation to the next in an organism.

### Persistent ancestral information modifies the cell code

A key concern in thinking about how organisms evolve is what changes in one generation can be passed on to subsequent generations. The nature of organisms presented here (Figure 1 and Figure 2) makes it clear that the persistence of changes across many generations requires modification of the cell code and thus evolution occurs through “descent with modification” (Darwin, 1859) of the cell code. Note that this does not include information (induction by signals, uptake of nucleic acids, etc.) accrued from additional sources such as the microbiome or maternal circulation during development that could also vary across generations.

Modifications to the cell code can alter its genetic (DNA) or non-genetic aspects (C_past_→n•E_past_→n). Changes to the DNA sequence can persist for many generations and are indeed the best-studied changes to the cell code. These can include both random mutations and changes derived from experience (e.g. CRISPR-Cas system of antiviral immunity in bacteria (Barrangou, 2015)). But, changes to the cell code that do not alter DNA sequence also have the potential to persist for many generations. In fact, one of the earliest mutants ever described - in 1744 - was found to not alter the DNA sequence. This variant of the toadflax *Linaria vulgaris* called *peloria* (greek pελωρ, monster) (Linnaeus and Rudberg, 1744; Gustafsson, 1979) arises at ∼1% frequency in each generation (DeVries, 1906) and is correlated with methylation but not sequence changes at the *Lcyc* locus (Cubas et al., 1999). Such inheritance of changes that do not alter DNA sequence and yet affect how the DNA sequence is used in any generation can be viewed as the influence of organismal history. The controlling influence of past cellular components on the production of future components from DNA is powerfully illustrated in ciliates that develop using RNA from one generation to splice or unscramble germline DNA sequences in the next generation. In these ciliates such as *Oxytricha* and *Tetrahymena*, changes in RNA introduced in one generation can alter how the DNA is rearranged in many subsequent generations (Bracht et al., 2013; Chalker et al., 2013). In other organisms, extensively studied phenomena like paramutation (Brink, 1956; reviewed in Hollick, 2017) and RNA interference (Fire et al., 1998) that cause persistent silencing of genes also illustrate the persistence of non-genetic information. While the precise mechanisms for how the gene silencing information is transmitted across generations are still being worked out, persistent chromatin modifications and amplified RNAs have emerged as possibilities (reviewed in Rankin, 2015). Transmission for many generations is likely to require inheritance of molecules containing the information for silencing a gene for one or a few generations combined with periodic replication of molecules to reinforce this information. For example, an ancestral event such as exposure to double-stranded RNA during RNA interference could trigger production of chromatin modifications in each generation based on instructions held in RNA. All that is needed to make this information stable for many generations is transmission, even for just one generation, followed by reproduction in every generation. Crucially, the resulting permanent changes to the cell code do not involve changes in the DNA sequence. Other mechanisms for changing the cell code without mutating DNA include the transmission of prions (Halfmann and Lindquist, 2010), where alternative folding states of proteins that can template recreation of similar states are transmitted across generations.

When complex organisms, like humans, that can make tools and artifacts by interacting with both the environment and other organisms are considered, there is no end to the longevity of ancestral information. For example, consider the impact of ideas in books on the lives of people. The spread of such ideas within a culture (i.e. a meme (Dawkins, 1989)) can have a profound influence on humans. Thus, the evolution of such organisms is shaped by much more than their cells (Jablonka and Lamb, 2005). In fact, contributions by many biological mechanisms that transmit information across generations can be incorporated into a consideration of how organisms evolve (Rivoire and Leibler, 2014), if unconstrained by evidence.

Even if only the transmission of biological material is considered, the precise way in which information is transmitted across generations is unclear. While DNA in a zygote is obtained only from the immediate parents, the provenance of the information contained in the non-genetic material within a zygote is less clear. If life originated once on earth as suggested by the commonalities among extant organisms, then every zygote can be traced back to the first cell by descent. This initial cell must have accrued complexities that were transmitted from one generation to the next. Therefore, although it is possible that the information contained in the non-genetic components and their arrangement is also from the immediate parents, it is also conceivable that this information initially arose in an ancestral generation and has since been faithfully propagated. Nevertheless, unlike in the case of genetic changes, there have only been a few cases of clear evidence for non-genetic changes that persist for many generations (see Jablonka and Raz, 2009 for review of early work). This paucity could reflect limited experimentation, incompatibility of the changes with the perpetuation of life, or mechanisms that oppose such persistence.

### Forces that oppose variation preserve the cell code

The developmental program of an organism imposes constraints on non-genetic variations within the germline, which conveys the cell code from one zygote to the next. The development of an organism in a given environment follows a path that reflects the teleonomic constraints imposed by epigenetic control systems within cells and cell-cell interactions in that organism. This constrained path was referred to as “chreod” (“necessary path” from greek roots χρη, it is necessary, and óδOζ, path (Waddington, 1957)) and has been given modern form within the framework of dynamical systems theory (Pisco et al., 2016). Such developmental constraints have been recognized as one of the forces that oppose change in organisms during evolution (Gould and Eldredge, 1993).

Some studies show correlation of certain molecular changes in an animal with environmental or dietary changes in parents or recent ancestors (e.g. Rodgers et al., 2013; Rechavi et al., 2014; Sharma et al., 2016; Chen et al., 2016; Klosin et al., 2017) and molecular changes such as histone methylation that occur within germ cells can cause effects that persist for a few generations (e.g. Greer et al., 2011; Siklenka et al., 2015). These instances of heritable changes reflect non-genetic modifications of the cell code and urge consideration of the possible ancestral origins of disease and the impact of medical intervention on descendants. However, loss or erasure of such chemical changes after a few generations reflects a homeostatic return to the original cell code. For example, during early mammalian development most parental chemical modifications such as DNA methylation are erased and new modifications are added (Feng et al., 2010). Because this happens in every generation, the information for adding these new modifications must be passed from one generation to the next as an aspect (molecule and/or arrangement) of the cell code. This developmental reprogramming thus preserves the cell code and opposes transgenerational epigenetic changes. Nevertheless, cases of persistent non-genetic changes - some lasting for tens to hundreds of generations (e.g., Vastenhouw et al., 2006; Ashe et al., 2012; Buckley et al., 2012; Luteijn et al., 2012; Shirayama et al., 2012; de Vanssay et al., 2012; Devanapally et al., 2015; Leopold et al., 2015; Minkina and Hunter, 2017; Lev et al., 2017; Ciabrelli et al., 2017; Devanapally et al., 2017) - provide us with opportunities to analyze non-genetic aspects of the cell code (Figure 3).

**Figure 3.**
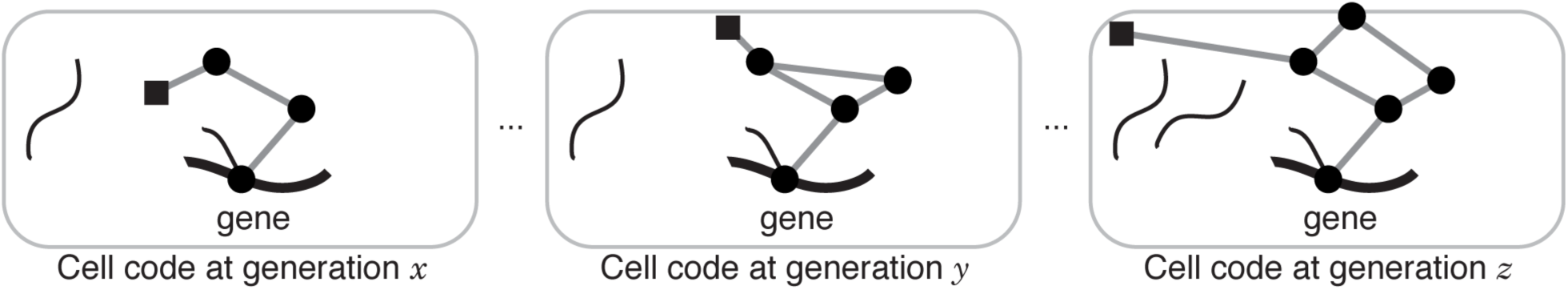
Ancestral non-genetic information and the evolution of the cell code. Schematic depicting possible evolution through changes to non-genetic components of the cell code from the perspective of a single gene. Past non-genetic changes (in generations *x, y, z*, say) impact the evolving cell code if they persist through life cycles and overcome homeostatic mechanisms that oppose changes to the cell code, information for adding these new modifications must be passed from one generation to the next as an aspect (molecule and/or arrangement) of the cell code. This developmental reprogramming thus preserves the cell code and opposes transgenerational epigenetic changes. Nevertheless, cases of persistent non-genetic changes - some lasting for tens to hundreds of generations (e.g., Vastenhouw et al., 2006; Ashe et al., 2012; Buckley et al., 2012;

Together, these considerations inform attempts to identify past events that have shaped the evolution of organisms and future attempts to synthesize organisms.

### To make life we need to know the cell code

Considering what we would need to know to make an organism from raw materials reveals the extent of our ignorance of the cell code. In recent years, we have begun to remove, replace, augment, or modify aspects of the cell code. In particular, techniques for manipulating DNA have advanced to the point that practically any change in DNA sequence can be made using genome engineering (reviewed in Komor et al., 2017) and even the entire genome of a simple organism can be replaced with a synthetic genome (Gibson et al., 2010). We can effectively transfer genetic material to avoid mutant mitochondria (Tachibana et al., 2009), we can augment the genetic code (Ostrov et al., 2016), and even use gene drives to change wild populations (Gantz and Bier, 2015). All these efforts are akin to modifying a machine by tinkering with or replacing parts in a pre-existing machine without fully understanding how the machine is put together. As with complex machinery, such manipulations without deep understanding can be dangerous and additionally all manipulations of life warrant careful ethical considerations (e.g. Baltimore et al., 2015). To make a machine from raw materials, however, we need to know the *entire* design. Because organisms build themselves, assembling the cell code of an organism could be sufficient to make an organism. Although an assembly of chemicals that display characteristics of life has been imagined in theory (e.g. The Chemoton Theory, Ganti, 2003), the enormity of this challenge in practice is clear in our struggle to use raw materials to make a rudimentary cell that perpetuates itself (Blain and Szostak, 2014; Schwille, 2015). Nevertheless, we have begun to coax pre-existing cells into making complex parts of organisms in vitro – e.g. eye (Eiraku et al., 2011) and brain (Lancaster et al., 2013). Eventually making entire complex organisms that reproduce will require first discovering the cell code of a simple organism and then determining the simplest version of that code. The “endless forms most beautiful and most wonderful” (Darwin, 1859) found in nature each have a different cell code. Such organisms sculpted by evolution, however, are unlikely to already have the minimal essential cell code because evolution tinkers with what exists (Jacob, 1977) but anticipates poorly. Our current understanding suggests that evolution proceeds through measure and counter measure, including nonadaptive processes (e.g. Lynch, 2007; Madhani, 2013), resulting in a cellular architecture that is more Rococo than Bauhaus with aspects that are superfluous (Gould and Lewontin, 1979). Thus, organisms found in nature have layers of post hoc regulation that can obscure essential design principles. Understanding such principles can enable the design of organisms free of the historically contingent tinkering of evolution.

### Approaches to decipher the cell code

Discovering the cell code of an organism minimally requires comparison of features in zygotes of successive generations (Figure 2). Our increasing ability to perform molecular studies on single cells (Tsioris et al., 2014) makes this approach reasonable. Parental effect mutants can be used to identify potential components of the cell code that are maternally or paternally deposited into the zygote. Such mutants may also identify signaling from one generation to the next. However, the arrangement of these molecules in the zygote is more difficult to discover and would require systematic cell biological and biochemical analyses. After these components are identified and their arrangements are discovered, the zygotes of successive generations can be compared.

Penetrating insights often require perturbation of the system and not mere observation. Past perturbation studies of development and reproduction have largely focused on the analysis of essential genes that impact viability or fertility, respectively. Consequently, the effects of a perturbation are typically only analyzed during a limited period of the life cycle. Furthermore, defects in an essential gene (required for development, say) would kill the organism precluding examination of the same stage in successive generations. Thus, when we interfere with an essential gene or process in one generation, we lose the ability to see its impact on the next generation because the intervening organism is affected. In other words, we cannot know if an essential gene or process is required to make the organism or to reproduce the cell code or both. Studies using viable and fertile mutants on the other hand permit examination of the zygote in successive generations. But, careful subsequent analysis will be needed to separate defects in the mere making of the organism from perturbations of the cell code. For example, a mutation in DNA that changes a residue in hemoglobin alters genome sequence in the cell code and affects the structure of the protein made in red blood cells, but likely does not affect non-genetic aspects of the cell code or its propagation via the germline. However, a mutation in DNA that changes a protein that is reproducibly present in the zygote or the germline changes not only the genome sequence in the cell code but potentially also non-genetic aspects of the cell code and/or its propagation. Finally, a non-genetic change (e.g. chromatin modification or addition of double-stranded RNA) that causes gene silencing that persists for many generations alters non-genetic aspects of the cell code and potentially its propagation without changing the genome sequence.

The ability to make random changes in DNA and examine its consequences – forward genetics – and more recently to turn off any gene through RNA interference – reverse genetics – were crucial for correlating changes in the genome with changes in the organism. To correlate changes in non-genetic components of the cell code with changes in the organism, additionally we need forward epigenetics and reverse epigenetics. While we do not yet have a way to perform forward epigenetics, reverse epigenetics has begun (Park et al., 2016). For example, a guide RNA and the Cas9 enzyme fused to a histone modifier can be used to target chromatin modifications to histones located in a specific region of the genome (e.g. Hilton et al., 2015). Such a manipulation could be performed in one generation and its impact analyzed in subsequent generations to determine whether the manipulation altered non-genetic aspects of the cell code.

Nuclear transplantation experiments were essential for the realization that most cells in an organism retain the same genome (Briggs and King, 1952; Gurdon et al., 1958). A similar approach could be used to discover all the cells that retain the ability to use the DNA genome to generate the entire organism, i.e., to discover all the cells that have the non-genetic aspects (molecules and arrangement) of the cell code. Every cell that can be induced to generate the entire organism is expected to either have, or be able to reconfigure its contents to create, the entire cell code. For sexually reproducing organisms, minimally every cell that is continuous within the female germline from one zygote to the next is expected to have all the non-genetic aspects of the cell code except those obtained from the male gamete upon fertilization. Conversely, every cell that is continuous within the male germline from one zygote to the next is expected to have all the non-genetic aspects of the cell code except those obtained from the female gamete upon fertilization. It will be interesting to discover the somatic cells that retain non-genetic aspects of the cell code despite differentiation during normal development and to discover how the cell code is reduced from being present in its entirety in the zygote to the portion in each gamete awaiting union upon fertilization.

In organisms that have well-differentiated somatic and germ cells (Extavour and Akam, 2003), it will be interesting to discover the extent of somatic influence on the perpetuation of the cell code via the germline. A germ cell from one animal could be transplanted into another (Brinster and Zimmermann, 1994) to evaluate the extent of somatic influence on the cell code. Similarly, the circulatory system of two animals could be joined together (reviewed in Eggel and Wyss-Coray, 2014) to examine the effect of secreted material from the cells of one animal on the germline of another. Evidence for such influence of secreted material is provided by intriguing observations like the entry of double-stranded RNA from neurons into the germline in the worm *C. elegans*, resulting in gene silencing that persists for more than 25 generations (Devanapally et al., 2015). Analyzing somatic influences in different environments could reveal mechanisms, if any, by which the environment could alter the cell code in different organisms.

Simpler organisms likely have simpler cell codes. Single-celled organisms like the marine green alga *Ostreococcus tauri* (Derelle et al., 2006) or bacteria could be chosen as model systems to discover the simplest cell codes. Simpler still are organisms that result from efforts to generate cells that have a minimal genome (Hutchison et al., 2016; Glass et al., 2017). While the analysis of these organisms would reveal the logic of cell codes for single-celled organisms, it is conceivable that the need for differentiation in complex animals and plants results in fundamentally different strategies for the assembly and reproduction of their cell codes.

An alternative to perturbation studies that overcomes the impasse of lethality or sterility when the cell code is disturbed is the use of tracer studies. To illustrate this approach, imagine you wanted to discover the extent of the circulatory system of an organism. A perturbation study could involve making cuts throughout its body and examining if blood spurts out. A tracer study involves injecting an inert molecule that permeates the entire circulatory system and then imaging that molecule. Therefore, taking the tracer approach to examine the cell code of an organism, we could insert benign sequences (e.g. encoding a fluorescent protein) into its genome, add or remove various regulatory features, and examine if these changes have consequences that persist across generations.

To make rapid progress in understanding the principles of the cell code, we are likely to benefit from focus. Examples from the history of biology support this assertion: the discovery of the genetic code began with a ‘tracer’ study where one RNA sequence (poly-U) was used to synthesize one peptide sequence (poly-Phe) in an *in vitro* translation system (Nirenberg and Matthei, 1961) and the basic principles of gene regulation were worked out through sustained effort on a few genes – e.g., the *lac* operon (Jacob and Monod, 1961). Similarly, focusing on one or a few genes, their protein and/or RNA products, and the factors that impact their recreation in successive generations will allow us to ask specific questions and generate a preliminary understanding of the cell code. This focused approach could be an effective complement to comprehensive approaches that compare many components of zygotes in successive generations in response to experimental manipulation.

## Conclusion

The sequence of a genome can be used to identify an organism. However, the genome sequence is not sufficient information for making that organism. To make an organism we need to know its cell code, which is the evolving arrangement of molecules, including the genome, that is nearly reproduced in every generation. Building on more than a century of work in biology, we can now begin to decipher the cell code of an organism by analyzing single cells that start successive generations.

## Acknowledgements

The ideas presented here were honed through long conversations with my friend Katerina Ragkousi, through lectures to students of a course on Ignorance run by José Feijo, and through a faculty chalk talk to colleagues at the University of Maryland. I thank Steve Wolniak for initially pointing me to relevant books on cell biology; Karen Carleton, David Mosser, Zhongchi Liu, Caren Chang, Norma Andrews, José Feijo, Wenxia Song, Carlos Machado, Tom Kocher, Charles Delwiche, William Snell, Steve Wolniak, Sougata Roy, Geraldine Seydoux, Kevin O’Connell, Andy Golden, John Hanover, Victor Ambros, Marie-Anne Felix, and Pierre-Emmanuel Jabin for comments on the manuscript; all past and present members of the Jose lab for lively discussions; and my friend John Horner for early support when these ideas were in their infancy. I apologize to any colleagues whose work could not be highlighted in this short article.

